# Methylation mimic mutations of progesterone receptor AF1 impair gene-specific regulation through stabilized chromatin interactions

**DOI:** 10.1101/2025.10.23.684277

**Authors:** Pheck Khee Lau, Bernett Lee Teck Kwong, Shi Hao Lee, Chew Leng Lim, Qian Yee Woo, Amanda Rui En Woo, Jace Koh, Valerie C.L Lin

## Abstract

Progesterone receptor (PR) is a nuclear receptor that regulates gene transcription through recruiting coregulators and general transcription factors by activation functions AF1 and AF2. AF1 localizes to the non-conserved and disordered N terminal domain and is believed to facilitate tissue- and gene-specific activity. Our previous proteomic analysis identified three key functional residues (K464, K481 and R492) of AF1 that are monomethylated. Mutations of KKR to phenylalanine (FFF) that mimic methylation created hypoactive PR, whereas KKR/QQQ mutations generated hyperactive PR in gene reporter assays. The current study investigated specific involvement of AF1 in PR regulation of gene expression in breast cancer cells. AF1-FFF mutations attenuated progestin induced growth regulation, cell adhesion and apoptosis and AF1-QQQ mutation enhanced these effects. Genome-wide expression analysis showed attenuated gene regulation by AF1-FFF of two thirds of PR target genes including genes involved in hypoxia and TNFα signalling via NFKB. Unexpectedly, AF1-FFF mutations had little effect on ligand-independent gene regulation, suggesting distinct mechanisms of gene regulation by liganded and unliganded PR. Intriguingly, impaired activity of methylation mimic mutant PR-FFF is associated with higher enhancer binding peaks in ChIP-Seq analysis. This corresponds to a stronger association of AF1-FFF with SRC-1 and AF2 as we previously reported. We propose that methylation mimic AF1 mutant impairs regulation of a subset of genes through tighter coregulator binding which restricts the dynamics of the disassembly of transcription complex.

## Introduction

Progesterone is critical for female reproductive processes including embryo implantation, maintenance of pregnancy and mammary development [1]. In the endometrium, progesterone exerts coordinated antagonistic and synergistic effect with estrogen to prepare for embryo implantation. This involves inhibition of estrogen-induced luminal epithelial cell proliferation in the preimplantation window and stimulation of glandular secretion. Progesterone also plays an important role in curbing the proliferative effect of estrogen on the endometrial lining to prevent endometriosis and endometrial cancer [2]. In the mammary gland, progesterone is involved in mammary stem cell expansion in the adult and critical for mammary ductal branching and alveologenesis [3, 4]. The role of progesterone in breast cancer has been controversial. Hormone replacement therapy (HRT) trials initially reported significant increased risk of breast cancer for women taking estrogen plus progestin compared to those taking estrogen alone [5, 6]. But these initial reports have been challenged, and it has been critically reviewed that there is no significant association between the use of progestin in HRT and breast cancer [7–10]. Consequently, there has been a revival of interest in progesterone receptor (PR) targeting therapies for breast cancer. Further understanding of the molecular mechanism of progesterone action is important for informed decision-making on PR targeted therapy.

PR is a direct estrogen-induced gene in most estrogen responsive cells and tissues, where PR expression is dependent on estrogen and usually is used as a marker of ERα activity in breast cancer. PR in turn acts synergistically or antagonistically with ER to regulate gene transcription [11–13]. Activated PR can redirect ER binding by modifying chromosome accessibility or recruit corepressor, alter the transcriptional activity or repress ER target genes [14]. PR exists in two main isoforms, PRA and PRB that are transcribed from different promoters resulting an additional 164 amino acids in PRB at its N terminus. PR is composed of a modular domain structure includes an N-terminal domain, (NTD), DNA-binding domain (DBD), hinge region (H), and hormone-binding domain (HBD) [15]. Like all NRs, PR activity is primarily mediated by Activation Function AF1 and AF2. PRB has an additional AF3 at its amino terminus, conferring its greater transcriptional activity compared to PRA. AF1 is located within the intrinsically disordered NTD and was found initially to be a weak ligand-independent activation domain at the time of discovery [16], while AF2 is located to the highly conserved and structured ligand binding domain (LBD) [17]. AF1 and AF2 synergizes to mediate ligand-induced transcriptional activity through recruiting transcription coregulators, such as the p160 family of coactivators SRC-1, SRC-2 and SRC-3, and corepressors such as NCoR1 and SMRT, which serve to recruit chromatin remodelers and the general transcriptional machinery [18, 19]. Dynamics of interaction between AF1/AF2 and coregulators can therefore influence gene regulation.

The non-conserved and intrinsically disordered nature of AF1 renders structural diversity and plasticity allowing malleable interaction and partner protein induced folding. For example, TATA-binding protein (TBP) binding to glucocorticoid receptor AF1 induces AF1 folding and facilitates its interaction with SRC-1 [20]. This structural plasticity and flexible interactions are said to be the basis for its cell and gene specific activities. However, the extent of AF1’s involvement in the cell- and gene-specific activity is still poorly understood. Unlike the highly conserved AF2, the primary sequence of AF1 is highly diverse among NRs, making it challenging to identify critical functional residues. We did proteomic profiling of post translational modifications of PR from T47D cells and identified 3 monomethylation sites, K464, K481 and R492 on AF1. Mutagenesis analysis showed that K464, K481 and R492 function cooperatively [21, 22]. Methylation mimic mutations KKR→FFF largely abolished PR activity while KKR→QQQ render PR hyperactive in reporter gene assays. Unexpectedly, the loss of activity of FFF is associated with stronger interaction with SRC-1 and AF2, and this may restrict the dynamics of the assembly and disassembly of transcription complex. Mice with homozygous FFF mutation are infertile due to implantation and decidualization defect [23]. These mice also exhibit defective mammary alveologenesis associated with low RANKL expression [24].

Targeting the AF1 of steroid hormone receptors is a promising strategy for cancer treatment. AF1 of androgen receptor has been extensively studied for targeting with small molecule inhibitors in prostate cancer [25–27]. A deeper insight into regulatory functions of PR AF1 in breast cancer is important for advancing PR-targeted therapies. The objectives of the current study were to elucidate the role of PR AF1 in the regulation of breast cancer cell growth and apoptosis, and to determine its extent of involvement in PR regulation of genome-wide gene expression and PR-chromatin interaction. In addition, the study evaluated whether AF1 is important in the synergistic and antagonistic interplay between PR and ERα in the context of whole genome regulation of gene expression.

## Results

### Stable expression of PR and AF1 mutants through lentiviral transduction

We reported that PR-FFF and PR-QQQ exhibited hypoactivity and hyperactivity in reporter gene assay, respectively [22]. The current study evaluated whether the activity of AF1 is gene and pathway specific in breast cancer cells, and whether AF1 is involved in PR-chromatin interactions. Stable expression of wild type PRB, PRB-FFF and PRB-QQQ were established through lentiviral transduction in MCF-7 cells, and all cells stably transduced with PR or mutant cDNA were selected in hygromycin containing medium and propagated into cell lines. Cells transduced with the empty viral vector (EV) were established as transfection controls. AF1 mutants exhibits normal nuclear localization characteristics (Supplementary Fig. 1). The immunoblot in Fig. 1A shows that the main PR expressed is PRB. Protein levels of PRB-FFF were slightly lower than PRB and PRB-QQQ. This is possibly due to greater proportion of PRB-FFF associated with chromatin as will be shown in ChIP-Seq data later. R5020 downregulated the level of ERα in PRB cells, and this effect was lost in PRB-FFF cells but remained in PRB-QQQ cells.

**Fig. 1.**
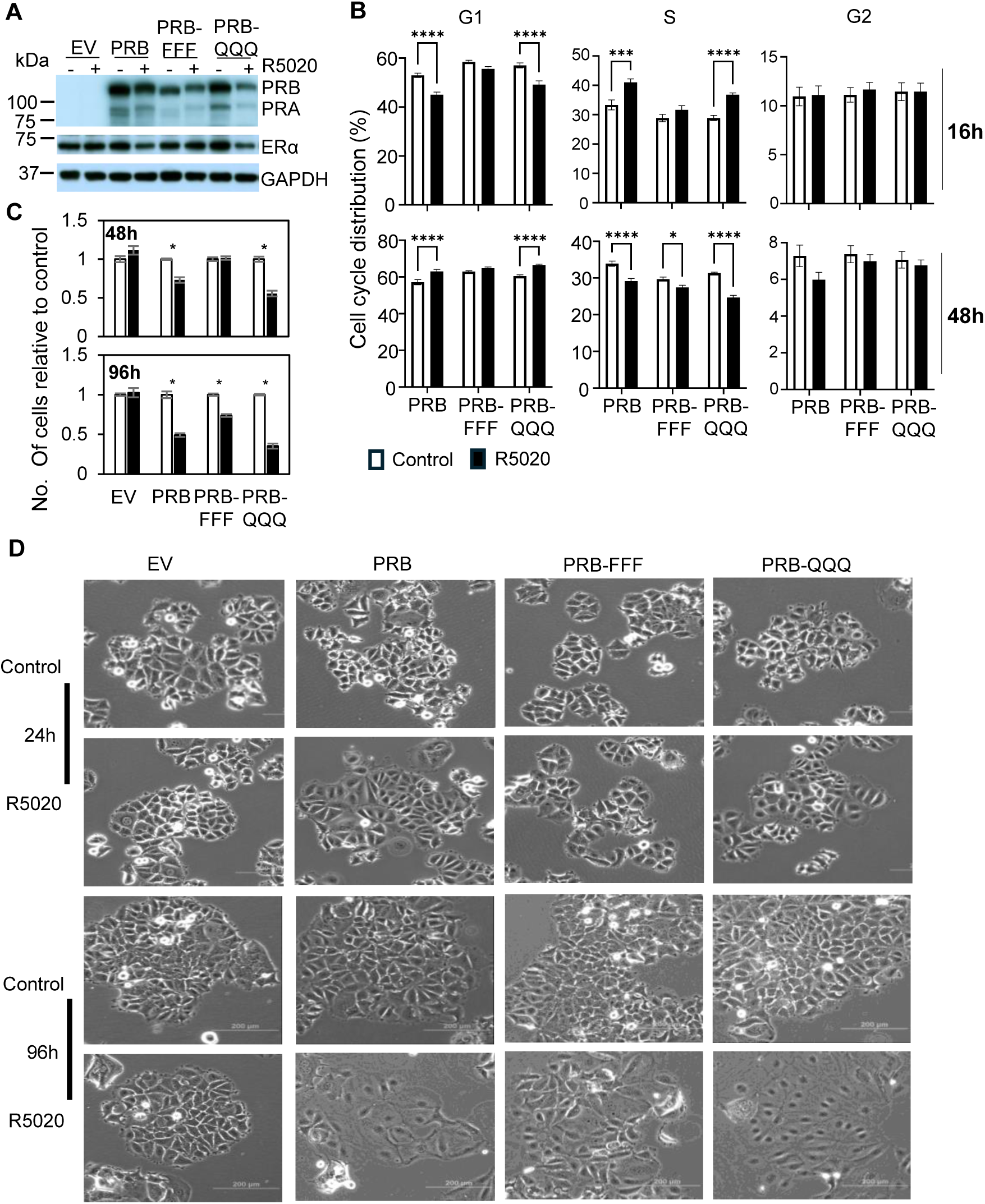
AF1 is important for PR to regulate growth and cell adhesion. **(A)** Western blotting analysis of PR and ERα protein levels in MCFD7 cells transduced with PRB and AF1 mutant cDNA. GAPDH was probed as a loading control. **(B)** Effect of R5020 on cell cycle progression measured by flow cytometry. Percentages of cells in G1, S and G2/M phase after 16h and 48h of R5020 treatment from four independent experiments (n=12) are presented. **(C)** Cell number after 48h and 96h of R5020 treatment in PRB, PRB-FFF and PRB-QQQ cells. Empty vector transduced cells (EV) served as controls. The data are presented as mean ± SEM (n = 6) and are expressed relative to the vehicle-treated cells which is set the value of 1. **(D)** R5020 induced cell spreading in PRB and PRB-QQQ cells after 24h treatment. PRB-FFF cells only exhibited cell spreading at 96 h postDtreatment. EV cells served as control which showed no response to R5020 treatment.

### AF1 is important for PR to regulate growth and cell adhesion

Progestin R5020 is known to exert biphasic effect on the cell cycle progression of breast cancer cells, in which R5020 accelerates G0/G1 to S phase progression initially, followed by G0/G1 arrest 24 – 36 hours later [28–30]. Consistently, R5020 reduced G0/G1 phase cells and increased S phase cells after 16h treatment of PRB cells (Fig. 1B), indicating that R5020 promoted cell cycle acceleration. After 48h treatment, R5020 increased fraction of G0/G1 cells, and reduced S phase fraction, indicating cell cycle arrest at G0/G1 phase. PRB-QQQ exerted similar biphasic effect on cell cycle progression. However, R5020 did not accelerate G0/G1 progression to S phase after 16 hours and its effect on G0/G1 phase arrest after 48 hours was impaired in PRB-FFF cells compared to PRB cells.

The effect of R5020 on cell proliferation was determined by cell counting with a hemocytometer. R5020 reduced cell number of PRB cells by 25% and 50% after 48h and 96h treatment, respectively (Fig. 1C). PRB-QQQ cells showed hyperactivity by reducing cell number by 40% and 65%, respectively. In contrast, R5020 treatment for 48h did not reduce cell number in PRB-FFF cells, and it reduced cell number by only 25% after 96h treatment, confirming the importance of AF1 in growth regulation in response to progestin.

AF1 mutations also altered effect of progestin on the cell morphology. R5020-induced growth inhibition was associated with cell spreading in PRB and PRB-QQQ cells after 24h treatment. R5020 induced cell spreading was more pronounced after 96h treatment (Fig. 1D). In contrast, R5020 induced cell spreading in PRB-FFF cells was not evident after 24h treatment and the cells were visibly less spread than PRB and PRB-QQQ cells after 96h treatment. Thus, AF1 is also involved in progestin induced signalling to cell adhesion.

### AF1 is important for PR to induce apoptosis in response to R5020

We reported that 96h treatment with R5020 induces marked apoptosis in MCF-7 cells transfected with PRB cDNA [51]. R5020 also caused marked apoptosis in PRB, PRB-FFF and PRB-QQQ cells after 96h treatment. But the magnitude of apoptosis, particularly late apoptosis was attenuated in PRB-FFF cells (Fig. 2A). For 72+48h time point, cells were sub-cultured after 72h treatment and treated for another 48h. There was significant induction of early apoptosis in all cell lines, but the magnitude of apoptosis was less in PRB-FFF cells. There was no significant induction of late apoptotic cells at 72+48h time point because these cells were only treated for 48 hours after subculture.

**Fig. 2.**
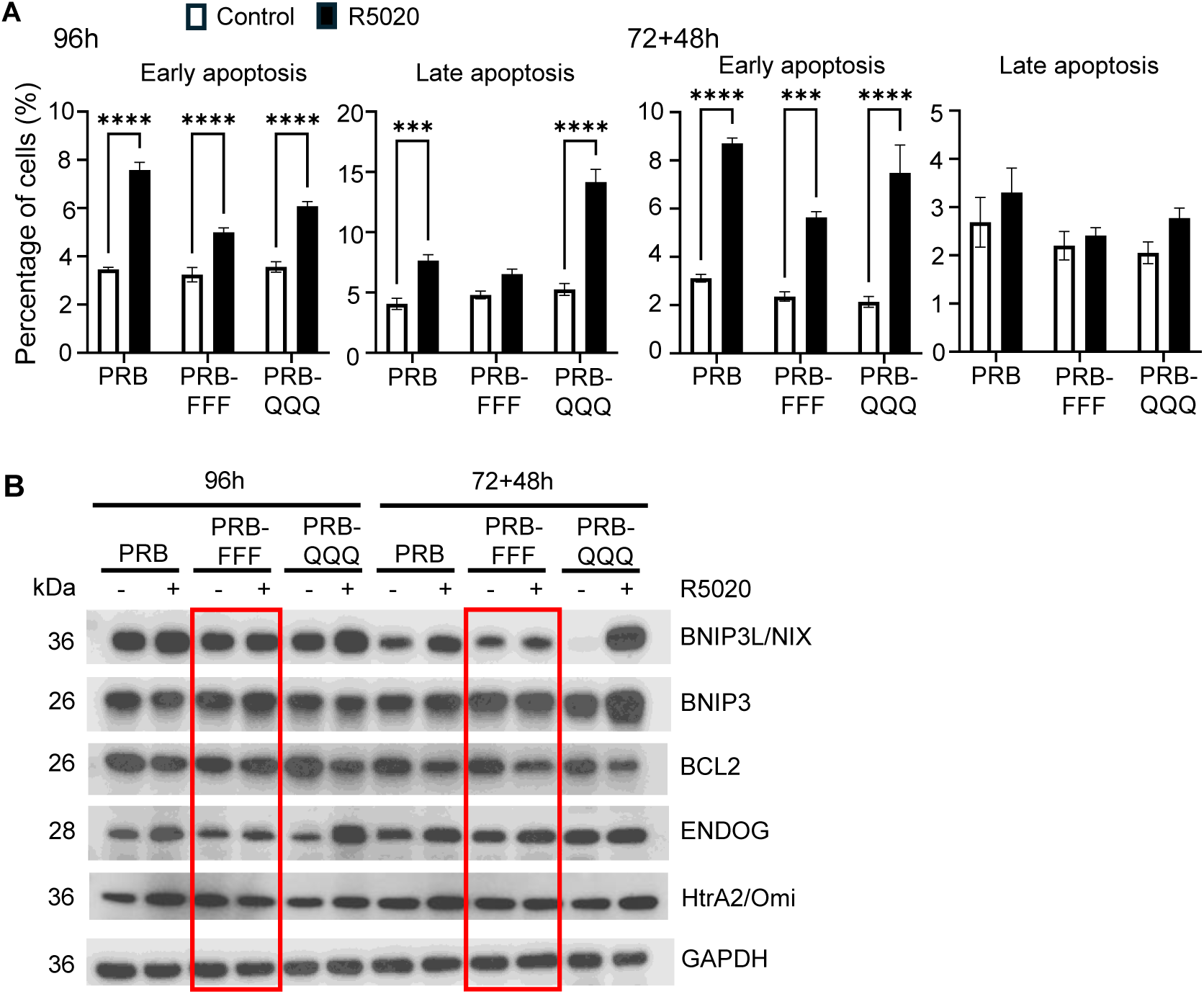
AF1 is important for PR to induce apoptosis in response to R5020. **(A)** Percentage of early and late apoptotic cells after 96h and 72+48h R5020 treatment in PRB, PRB-FFF and PRB-QQQ cells are presented (n=6 from two independent experiments). Apoptosis was attenuated in PRB-FFF cells. **(B)** Anti-apoptotic protein and mitochondrial pro-apoptotic proteins were analyzed by Western blotting analysis. R5020-mediated regulation in PRB-FFF cells was impaired (in red box). GAPDH was probed as a loading control.

Previous proteomic analysis showed that R5020 induced mitochondria mediated cell death in MCF-7PRB cells through upregulating mitochondria proapoptotic factors (https://www.biorxiv.org/content/10.1101/2025.03.06.641096v1.full). Western blotting analysis revealed that BNIP3 and NIX/BNIP3L, which are the pro-apoptotic members of Bcl-2 family, were upregulated in PRB and PRB-QQQ cells after 96h and 72+48h R5020 treatment, while anti-apoptotic protein Bcl-2 was downregulated. (Fig. 2B). Additionally, mitochondrial pro-apoptotic protein HtrA2/Omi and ENDOG were upregulated at both timepoints for PRB and PRB-QQQ cells. HtrA2/Omi is a serine protease released from the mitochondria and promotes cell death by inhibiting apoptosis inhibitors [31]. ENDOG is a DNA nuclease released from the mitochondria in response to apoptosis signals and induces DNA degradation in the nucleus [32]. Consistent with its impaired apoptotic response, the regulation of these proteins was substantially attenuated in PRB-FFF cells. Together, these function studies indicate that AF1 is involved in PR regulation of cell growth, adhesion and apoptosis.

### PRB-FFF exhibits hypoactivity in regulation of about two thirds of PR target genes

#### Hypoactivity of PRB-FFF and hyperactivity of PRB-QQQ in subset of PR target genes

To investigate genome-wide activity of AF1, bulk RNA-Seq analysis was conducted for PRB, PRB-FFF and PRB-QQQ cells in response to 6h R5020 treatment. DESeq2 analysis revealed 1950, 1205, 3158 significantly regulated genes (padj <0.05) by PRB, PRB-FFF and PRB-QQQ, respectively. Volcano plots show that the R5020-induced fold changes and statistical significance in PRB-FFF cells are generally lower, while those in PRB-QQQ cells are generally higher than in PRB cells (Fig. 3A).

**Fig. 3.**
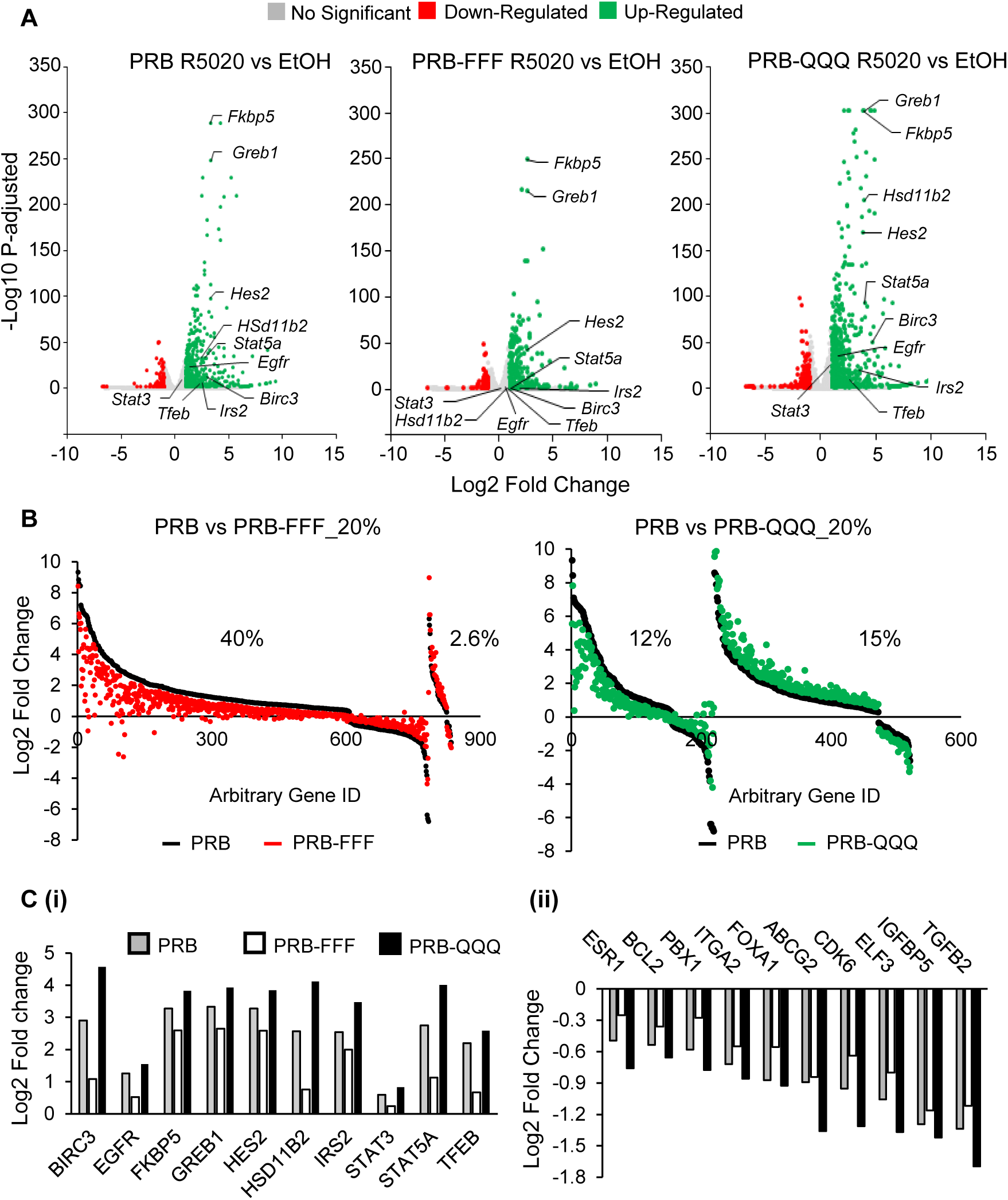
PRB-FFF cells exhibit hypoactivity while PRB-QQQ cells exhibit hyperactivity in gene regulation in response to R5020. **(A)** Volcano plots show significantly regulated genes in PRB, PRB-FFF and PRB-QQQ cells after 6h R5020 treatment. Ten well-known PR target genes are marked out. **(B)** PRB-FFF hypo-regulated nearly half of the PRB-regulated genes, whereas PRB-QQQ hyper-regulated ∼15% of the genes in response to 6h R5020 treatment. **(C)** PRB-FFF cells showed attenuated regulation of well-known PR target genes, regardless of whether they were upregulated or downregulated.

Further analysis shows that 40% of PRB regulated genes (pdaj <0.05) were hypo-regulated by PRB-FFF mutant by more than 20% (Fig. 3B), and 67% were hypo-regulated by more than 10% (not shown). Thus, KKR/FFF mutations hinder AF1 activity on two thirds of PR target genes. In contrast, 15% of the genes were hyper-regulated and 12% were hypo-regulated by QQQ mutant by more than 20%. These gene expression data is consistent with the hypo and hyper activity of PRB-FFF and PRB-QQQ in PR report gene assay [22], although the effect of QQQ mutations was more gene specific. Fig. 3C shows 10 well known up- or down-regulated PR target genes such as FKBP5, HSD11B2 and STAT5A that were hypo-regulated and hyper-regulated by PRB-FFF and PRB-QQQ, respectively (Fig. 3C).

#### AF1 is involved in most PR regulated pathways although the activity is gene specific

Gene set enrichment analysis (GSEA) was conducted to see whether these AF1 regulated genes belong to specific functional categories or in specific pathways. Positively enriched hallmark gensets with FDR <0.25 and normalized enrichment score >1.2 in all the three cell lines were listed in Fig. 4A. The common Hallmark gensets enriched by both PRB and PRB-QQQ cells treated with R5020 include ESTROGEN_RESPONSE_EARLY, ESTROGEN_RESPONSE_LATE, KRAS_SIGANLLING_DN, TNFA SIGNALING VIA NFKB, HYPOXIA and UNFOLDED_PROTEIN_RESPONSE. However, PRB-FFF regulated genes showed no significant enrichment for any of these Hallmark gene sets. For example, 85% of PR regulated genes (padj<0.05) in HYPOXIA (Fig. 4Bi) and TNFA SIGNALING VIA NFKB hallmark (Fig. 4Bii, Supplementary Doc 1) were hypo-regulated by PRB-FFF.

**Fig. 4.**
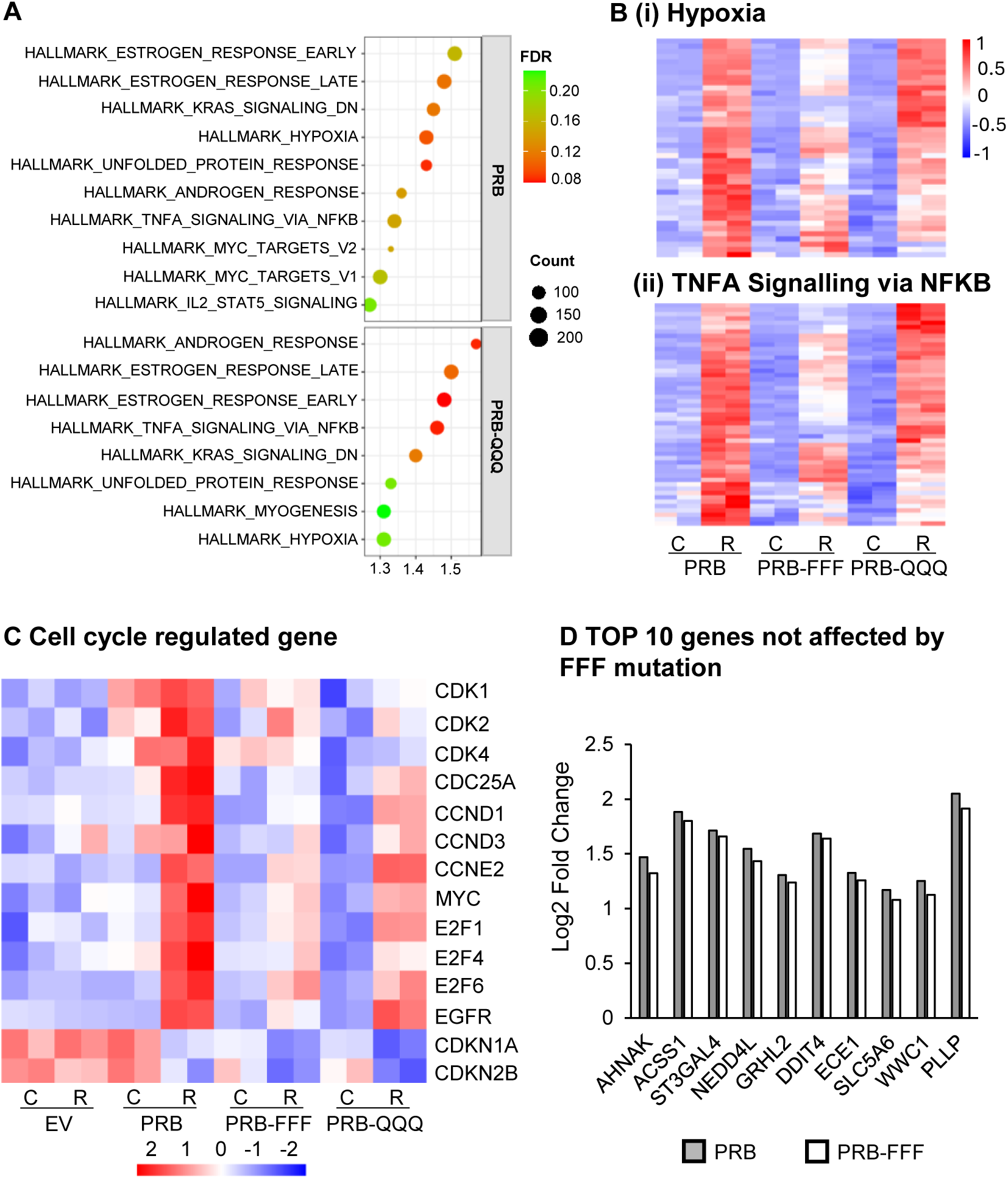
AF1 is involved in PR regulated pathways but is gene specific. **(A)** GSEA revealed no hallmark gene set was significantly enriched by R5020 in PRB-FFF cells. The bubble plot shows hallmark gene sets with FDR <0.25 and enrichment score > 1.2 enriched in PRB and PRB-QQQ cells. **(B)** Heatmap of genes associated with HALLMARK_HYPOXIA **(i)** and HALLMARK_TNFA_SIGNALING_VIA_NFKB **(ii)** indicates ∼85% of the genes in PRB-FFF cells are hypo-regulated relative to PRB, suggesting loss of function in response to R5020. C: Control; R: R5020. **(C)** Heatmap of cell cycle regulated genes shows attenuated regulation in PRB-FFF cells compared to PRB and PRB-QQQ cells, consistent with delayed cell cycle progression. C: Control; R: R5020. **(D)** Top 10 genes unaffected by FFF mutation in response to R5020 (based on padj value) are listed. The genes are involved in diverse molecular functions, indicating the effect of FFF mutation is gene specific.

Since PRB-FFF was unable to induce the initial phase of cell cycle acceleration in response to R5020, we asked whether this loss of activity is reflected at the level of gene regulation at the early time point. Fig. 4C shows that 6 hour’s treatment with R5020 upregulated the expression of cyclin dependent kinases (*Cdk1*, *Cdk2* and *Cdk4*), cyclins (*Ccnd1*, *Ccnd3* and *Ccne2*) and other growth promoting proteins such as E2F transcription factors, *Egfr* and *Myc* in PRB cells. R5020 also downregulated CDK inhibitors, *Cdkn1a* and *Cdkn2b*. Whereas these genes were similarly upregulated in PRB-QQQ cells in terms of fold changes, the effect of R5020 on these genes was much attenuated in PRB-FFF cells (Fig. 4C)

On the other hand, the expression of 541 genes out of 1950 R5020-regulated genes (∼28%) were not affected by AF1-FFF mutations by more than 10% (Supplementary Doc 2), indicating that AF1 is not required for PR to regulate these genes. However, GSEA or GO analysis did not yield significant enrichment of these AF1-independent genes in pathways or functional categories, indicating these genes are involved in diverse molecular functions. For example, the function of top 10 progesterone-regulated genes in this list (Fig. 4D) include cell adhesion and calcium signalling (*Ahnak*), regulation of cell adhesion and morphogenesis, (*Grhl2*), metabolism and bioenergetics (*Acss1*, *Ddit4*, *Slc5a6*), and protein ubiquitination (*Nedd4l*). Taken together, AF1 is important for the regulation of two thirds of PR target genes.

### Effect of AF1 mutations on ligand independent gene regulation is distinct from ligand-dependent gene regulation

AF1 was initially discovered as a ligand-independent activation domain in gene reporter assays and studies have shown that PR can regulate gene expression in a ligand independent manner through phosphorylation by growth factor-initiated signalling [33–35]. Comparison between vector-transfected cells (EV cells) and PRB cells revealed 232 differentially expressed genes in the absence of R5020 (p<0.01) (Fig. 5A, Supplementary Doc. 3). Interestingly, QQQ mutations abolished the upregulation of most genes by unliganded PRB, but FFF mutations had effect on only a few genes such as *Ass1*, *Aldh3b2*, and *Krt17*. No notable effect of AF1 mutation was observed for the downregulated genes. Hence the effects of AF1 mutations on ligand-independent gene regulation are distinct from that on ligand-dependent gene regulation, in which PRB-FFF is generally hypoactive and PRB-QQQ is hyperactive. It is also interesting to note that these ligand independently regulated genes were not typically regulated by the ligand R5020. Only 18 of the 232 genes were upregulated by R5020 (Fig. 5B, boxed region), but these 18 genes were downregulated by unliganded PRB.

**Fig. 5.**
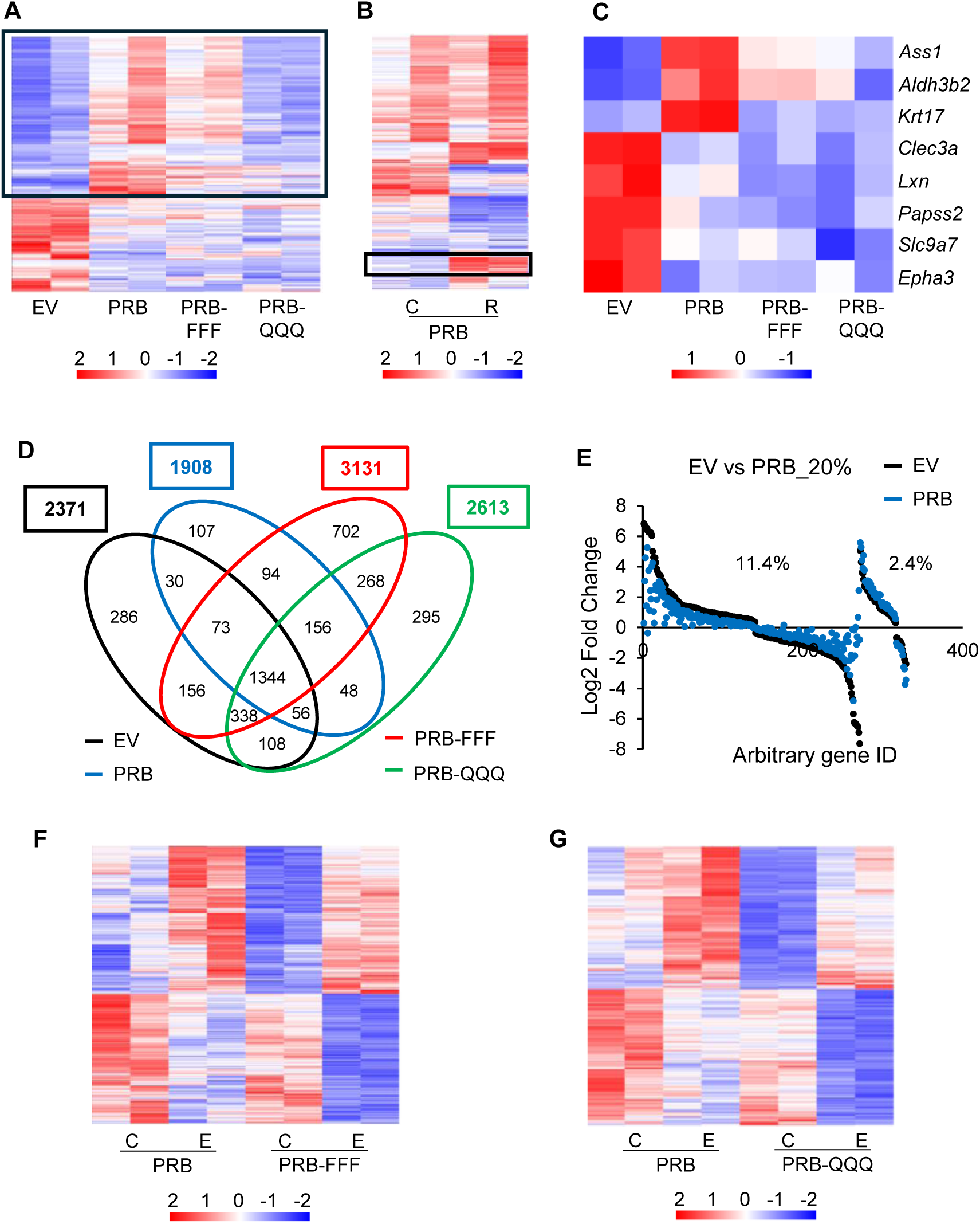
AF1 is required for ligand independent gene activation but not suppression by PR. **(A)** List of ligand independent PR-regulated genes (p < 0.01). Black box highlights reduced regulation by AF1 mutants compared to PRB. **(B)** Heatmap of ligand independent PR-regulated genes in PRB cells in response to R5020. C: Control; R: R5020. **(C)** List of ligand independent PR-regulated genes with padj < 0.05. **(D)** Venn diagram indicaties the overlapping of genes significantly regulated by E2 in EV, PRB, PRB-FFF and PRB-QQQ cells (padj < 0.05). **(E)** PRB exerts some repressive effects on as subset of estrogen regulated genes independent of R5020. **(F, G)** Comparison of unique E2-regulated gene in PRB-FFF cells **(F)** and in PRB-QQQ cells **(G)** with that in PRB cells shows that the basal levels of expression of these genes differ between AF1 mutants and PRB cells.

Functions of the unliganded PR target genes are varied. Fig. 5C shows top 8 genes regulated by unliganded PR based on padj <0.05. There top 3 upregulated genes are *Ass1*, *Aldh3b2* and *Krt17*. *Ass1* (Argininosuccinate synthetase) catalyses the formation of argininosuccinate from citrulline and aspartate, a key step in the urea cycle and in the synthesis of arginine. Ass1 is upregulated in response to DNA damage and halts cell-cycle progression by limiting nucleotide synthesis and gene transcription of a subset of p53 regulated genes [36]. It is also a tumor suppressor gene in triple negative breast cancer [37]. *Aldh3b2* is an aldehyde dehydrogenase for aldehyde detoxification and has been reported to promote the growth and invasion of cholangiocarcinoma [38]. *Krt17* is a component of the intermediate filament, and its high expression is associated with idiopathic pulmonary fibrosis [39]. Notably, ligand-independent upregulation of these three genes is AF1 dependent as AF1-FFF or AF1-QQQ mutations weakened the upregulations.

We also investigated whether PR exerts ligand-independent effect on estrogen regulated gene expression, and whether the effect involves the activity of AF1. After treatment with control vehicle or E2 for 6h, 2391, 1908, 3131 and 2613 genes were significantly regulated in EV, PRB, PRB-FFF and PRB-QQQ cells, respectively (padj < 0.05, Fig. 5D). 1344 genes were significantly regulated in all the 4 cell lines, while 286 genes, 107 genes, 702 genes and 295 genes were uniquely regulated significantly by EV, PRB, PRB-FFF and PRB-QQQ, respectively. E2 regulated fewer genes in PRB cells than in EV cells, with 11.4% (271) of the genes being less regulated by more than 20% in PRB cells based on fold change (Fig. 5E). Hence, PRB exerts some repressive effects on a subset of estrogen regulated genes independent of R5020. On the other and, AF1 mutations did not have definite effect on how PRB influences E2-induced gene expression despite of 40% and 27% more genes were significantly regulated by E2 in PRB-FFF and PRB-QQQ cells respectively (Fig. 5F, 5G). These genes were also regulated by PRB, albeit insignificant statistically due to higher or lower basal levels in untreated cells. It should be noted that each of these cell lines consists of all antibiotic-resistant cells after viral transduction. The differences in basal expression of some of the genes are not due to clonal selection. It is thus likely that AF1 mutants influence the expression of these genes in the absence of ligand.

### AF1 mutations impair synergistic and antagonistic activity between E2 and R5020 due to weakened response to R5020

Since progesterone is known to exert synergistic and antagonistic effects with estrogen on gene regulation, we investigated whether AF1 is involved in this functional interplay. DESEQ2 analysis showed that 2823 genes were significantly regulated in PRB cells in response to combined E2+R5020 treatment, as compared to 1908 genes in response to E2 and 1950 genes in response to R5020 alone (Supplementary Fig. 2A). 41% of E2-regulated genes in PRB cells were differentially regulated by greater than 20% in response to E2+R5020 (Supplementary Fig. 2B), suggesting co-regulation by E2 and R5020. PRB-FFF mutations attenuated the effect of R5020 on some of the E2 regulated genes, resulting mainly less up- or downregulated in between E2+R5020 and E2 treated PRB-FFF cells compared to PRB cells (Fig. 6Ai). On the other hand, fold changes between E2+R5020 and E2 treated PRB-QQQ cells can be more or less than that in PRB cells (Fig. 6Aii). Therefore, AF1 plays a role in the functional interplay between estrogen and progestin and FFF and QQQ mutations influence the interplay differently.

**Fig. 6.**
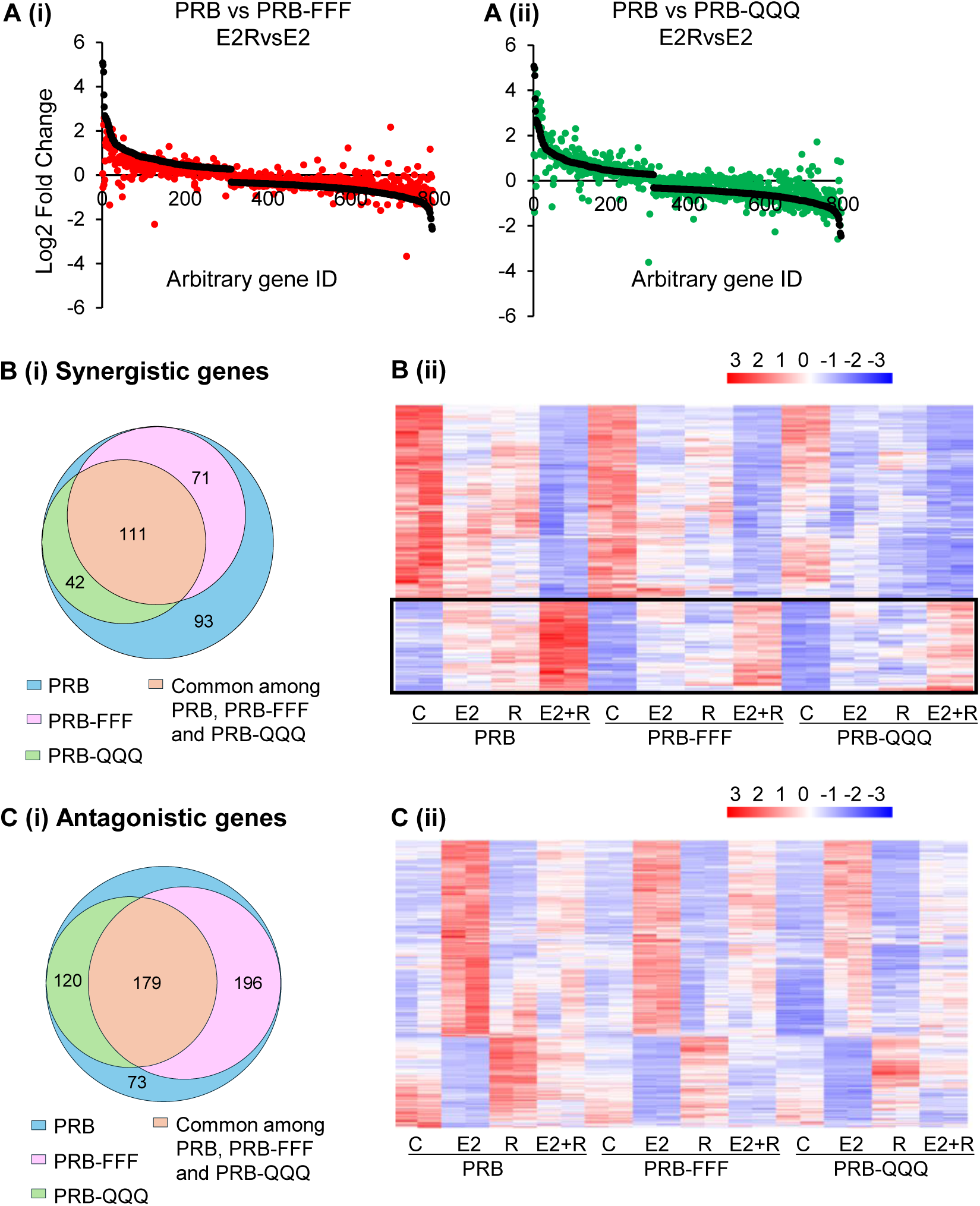
AF1 mutant impair synergistic or antagonistic activity of E2 and R5020. **(A)** The genes shows more than 20% difference in regulation between E2 and E2+R5020 treatment in PRB are listed and compared with PRB-FFF and PRB-QQQ cells. **(B)** Synergistic genes identified in PRB cells and the comparison with PRB-FFF and PRB-QQQ cells. **(i)** Venn diagram shows the overlap of synergistic genes between PRB, PRB-FFF and PRB-QQQ cells. **(ii)** Heatmap of synergistic genes. Boxed region highlights genes for which AF1 mutations attenuate the synergistic effect. C: Control; E2: Estradiol; R: R5020; E2+R: Estradiol and R5020. **(C)** Antagonistic genes identified in PRB cells and the comparison to PRB-FFF and PRB-QQQ cells. **(i)** Venn diagram indicates the number of overlapping antagonistic genes in PRB, PRB-FFF and PRB-QQQ cells. **(ii)** Heatmap of antagonistic genes. C: Control; E2: Estradiol; R: R5020; E2+R: Estradiol and R5020.

To define genes synergistically regulated by E2 and R5020, E2+R5020 regulated genes must be significant over vehicle treated, E2-treated and R5020-treated alone. Additionally, the magnitude of gene regulation by E2+R5020 must be greater by 20% than that by R5020 or E2 alone. This yielded 317 genes from PRB cells, with 102 upregulated genes and 215 downregulated genes (Supplementary Doc. 4). 93 genes were uniquely regulated by PRB, suggesting that AF1 mutations alter the regulation (Fig. 6Bi). Heatmap of the 317 genes shows that AF1 mutations attenuated the synergistic effect on upregulated genes (Fig. 6Bii, boxed region) largely due to less upregulation in response to R5020 or E2 in PRB-FFF and PRB-QQQ cells. On the other hand, AF1 mutations did not change the synergistic downregulation notably.

Antagonistically regulated genes are defined as E2-regulated genes whose magnitude of regulation by E2+R5020 were at least 20% less than that by E2 alone or exhibited an opposite regulatory direction. 568 of E2 target genes were thus listed as antagonistically regulated by R5020 in PRB cells (Supplementary Doc. 5). 73 genes were uniquely regulated by PRB (Figure 6Ci). Heatmap of the 568 genes shows that, for genes upregulated by E2, R5020 alone has no effect but antagonized E2-induced upregulation (Fig. 6Cii). AF1 mutations did not affect combined effect notably. For genes downregulated by E2 but upregulated by R5020, AF1 mutations impaired R5020-induced upregulation in most genes and the combined effects of E2 and R5020 are largely the average of the two effects.

### PRB-FFF exhibits greater Genome-wide binding than PRB and PRB-QQQ

#### ChIP-Seq analysis detected more PRB-FFF binding sites than PRB

We reported that FFF mutation enhanced the ligand-induced interaction between AF1/NTD and SRC-1, and between AF1/NTD and AF2/LBD [22]. We proposed a tripartite model in which the transient and calibrated interactions among AF1, AF2 and coregulators are important for a productive enhancer and dynamic cycling of the transcription complex. We hypothesize that more stable interaction due to PRB-FFF mutation results in prolonged enhancer occupancy and slower transcription rate. The prolonged enhancer occupancy would mean higher levels of PRB-FFF chromatin binding. Indeed, ChIP-qPCR showed that levels of PRB-FFF binding to the enhancer regions of FKBP5 and Chr 6p24.1-25.3 were higher than that of PRB and PRB-QQQ (Fig. 7A).

**Fig. 7.**
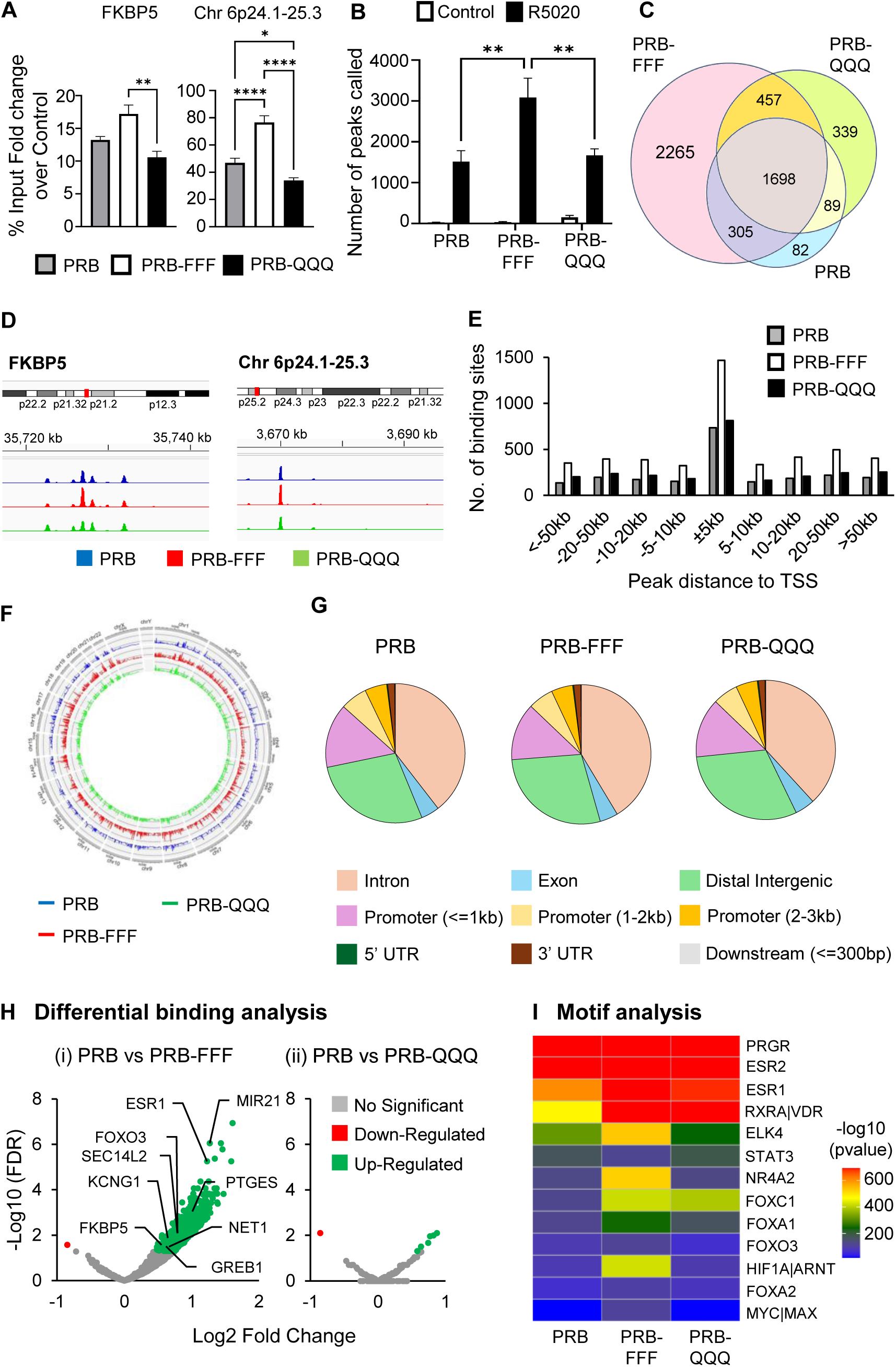
PRB-FFF exhibits greater genome-wide binding than PRB. **(A)** ChIP-qPCR analysis of PR target genes showing greater enhancer binding of PRB-FFF in response to R5020 treatment. Data are expressed as percentage of input, calculated as the ratio of signal for R5020 treated to vehicle treated cells. **(B)** Number of peaks called in PRB, PRB-FFF and PRB-QQQ cells after 1 hour of vehicle (ETOH) or R5020 treatment. Data in triplicates are plotted. **(C)** Venn diagram shows the overlap of R5020-induced peaks among PRB, PRB-FFF and PRB-QQQ cells. **(D)** Genome browser tracks illustrates the binding peaks at representative PR target genes. **(E)** Distribution of PR binding sites relative to their nearest TSS. Highest enrichment occurs at ±5kb of the TSS across all cell lines, with PRB-FFF exhibiting the highest number of binding sites. **(F)** Circle plot of PRB peak density across all chromosome regions. PRB-FFF exhibits enriched binding genome wide. **(G)** Distributions of PR peaks by PRB, PRB-FFF and PRB-QQQ in different gene regions are highly similar. **(H)** Volcano plots show significant enriched differential binding by PRB-FFF but not by PRB-QQQ compared to PRB **(I)** Relative enrichment of transcription factor binding motifs associated with PR peaks in PRB, PRB-FFF and PRB-QQQ cells. Negative logarithm of p-value is shown, blue indicates non-significant enrichment, and yellow to red indicates increasing significance.

To determine if higher PRB-FFF binding is a genome wide phenomenon, ChIP-Seq was conducted in triplicates. The number of peaks identified from each control and R5020 treated sample shows that the number of peaks called by R5020 treated PRB-FFF cells is on average more than 2 times of PRB peaks, indicating its stronger binding in target regions consistently (Fig. 7B). Based on pooled average peak sets, 2177 of PRB binding peaks were found for PRB. In contrast, 4725 binding peaks were identified for PRB-FFF and 2584 binding peaks for PRB-QQQ. 47% PRB-FFF binding overlaps with PRB and 69% PRB-QQQ binding overlaps with PRB (Fig. 7C). Although the number of PRB binding peaks is considerably lower compared to the reported number for T47D cells [40], the analysis confirmed PR binding to most previously known PR-target genes such as *Fkbp5* and *Chr 6p24.1-25.3* (Fig. 7D), *Hsd11b2*, *Net1* and *Stat5a* (Supplementary Fig. 3A). Furthermore, PR binding is approximately 5-fold higher within the 5kb of the TSS compared to other regions, regardless of the cell line (Fig. 7E). Consistently, higher PRB-FFF binding peaks are seen in regions from -5 kb to +5 kb (Fig. 7E). In addition, more PRB-FFF bindings are observed across all regions of chromosomes (Fig. 7F). On the other hand, binding distributions across promoter regions, introns, intergenic regions etc were similar between PRB and AF1 mutants (Fig. 7G).

#### PRB-FFF binding to transcription factor motifs is significantly higher than PRB

Diffbind analysis showed 902 differentially bound regions between R5020-treated PRB-FFF and PRB (FDR < 0.05), of which 901 peaks show stronger binding by PRB-FFF compared to PRB (Fig. 7Hi). The binding to known PR target genes *Esr1*, *Mir21*, *Foxo3* and *Ptegs* are among the genes with more PRB-FFF binding. In contrast, PRB-QQQ showed only a few genes with significant differential binding (Fig. 7Hii). Higher levels of chromatin binding by PRB-FFF are likely due to its stronger interaction with SRC-1 and AF2 [22], which could slow down transcription complex disassembly. Hence, the data support the notion that AF1 is involved in modulating dynamics of PR-chromatin interaction.

Expectedly, motif analysis using MEME Suite revealed that the topmost enriched binding motif of PRB, PRB-FFF and PRB-QQQ is PR consensus motif GGTACANNNT-GTTCT (Supplementary Fig. 3B). Other enriched motifs belong to zinc finger family of proteins. Because our dataset contained relatively low number of peaks, yielding only limited number of motifs, we employed another motif discovery tool MDSeqPos to further identify the motifs enriched. The peak regions were scanned, and the top 3 enriched motifs were for PGR, ESR1 and ESR2 binding (Fig. 7I). This is consistent with the understanding that PR is often associated with ESR and recruited to its binding sites in response to progestin [14]. In addition, PRB-FFF binding sites were also more enriched with motifs for ELK4, NR4A2, FOXA1 and HIF1α than PRB and PRB-QQQ. ELK4 is a ETS domain containing protein and is also reported to be enriched at binding sites for ERα/PR complexes [41]. *Nr4a2* is a nuclear orphan receptor and its known target gene *Bndf* is also a PR target gene in breast cancer cells and in neuronal tissue [42, 43]. *Foxa1* is pioneer factor that opens chromatin regions to allow for ERα and PR access [44]. Its binding sites are known to be overrepresented in PRB binding regions [45]. It is plausible that AF1 plays a role in modulating PR regulation of gene expression with *Elk4*, *Nr4a2*, *Foxa1* and *Hif1*_α_.

## Discussion

We previously reported important roles of PR AF1 in endometrial and mammary development in mouse models with AF1-FFF and AF1-QQQ mutants. The current study evaluated roles of AF1 in progestin regulation of cellular activity and gene expression in breast cancer cells. The study made three lines of significant findings. First, AF1 is involved in progestin regulation of cell growth, adhesion and apoptosis, and PRB-FFF mutant exhibited significant impairment in these activities. Second, AF1 activity is gene specific. AF1 mutations affected regulation of approximately two thirds of PR target genes by progestin. Furthermore, ligand-dependent and independent activities of AF1 are mediated through distinct mechanisms because FFF and QQQ mutations affected ligand-dependent and independent activities very differently. Importantly, PR AF1 is involved in modulating PR-chromatin interaction as the hypoactive mutant PRB-FFF shows more enhancer binding than PRB, likely due to stronger interaction with SRC-1 and AF2 [22]. Taken together, PR AF1 is important for the core activities of liganded PR in regulating growth and it is involved in the modulation of PR-chromatin interactions although structural plasticity of AF1 likely allows it to contribute to gene regulation through multifaceted mechanisms.

AF1 was originally discovered as a ligand-independent activation domain of chick PR using reporter gene assays [16]. Ligand-independent activity of PR and steroid hormone receptors is traditionally defined by activity in reporter gene assay, a phenotype such as growth, or interaction with transcription coactivators [46, 47]. It may be influenced by the cellular context, or expression of coregulators such as PRMT6 [48]. One unexpected finding from our RNA-Seq analysis is that genes regulated by unliganded PR are either not regulated by R5020, or for a few genes regulated by R5020 in the opposite direction of unliganded PR. This suggests that the mechanism of ligand-independent activity of PR is different from the ligand induced activity. Second unexpected finding is that property of amino acid for ligand-induced activity of AF1 is different from the ligand-independent activity. The hyperactive PRB-QQQ in the presence of ligand becomes hypoactive in gene activation by unliganded PRB, whereas PRB-FFF had little effect. It is plausible that the ligand-independent activity of PR in gene activation involves AF1 and KKR methylation because FFF mutant with increased local bulkiness and hydrophobicity works well for the ligand-independent gene activation.

The dynamics of nuclear receptor-chromatin interactions has direct impact on transcription rate and physiological outcomes [49, 50]. It is influenced by binding dynamics with coregulators, chaperones, general transcription factors as well as chromatin structure [51–53]. We propose that PR AF1 is involved in modulating the dynamics of PR-chromatin interaction through its interaction with coregulators and the general transcription machinery. This is based on the following evidence. First, loss of function mutant PRB-FFF exhibits significantly stronger interaction with SRC-1 and AF2, and this is associated with markedly slower receptor activation kinetics of the mutant [22]. Second, these stronger interactions are associated with more chromatin occupancy as indicated by more binding of PRB-FFF mutant than the PRB in ChIP-Seq analysis. More chromatin binding is likely due to a slower rate of dissociation resulting from more stable binding with its partners and hence slower activation kinetics. Additionally, modulation of PR-chromatin interaction by AF1 could be regulated by post translational modifications. We have shown that K464, K481 and R492 are monomethylated [22, 54]. The bulky and hydrophobic phenylalanine (F) can mimic the increased bulkiness and hydrophobicity of methylation. Since AF1 does not binds DNA directly, monomethylation of AF1 could stabilize its interaction with partners such as SRC-1 and AF2 and this may impair disassembly kinetics of PR transcription complex.

p160 family coactivators SRC1, SRC2 and SRC3 are common partners for AF1. PR Cryo-EM structure shows direct interaction of AF1 with SRC-2 [55]. AF1 also binds to TATA box binding protein (TBP) and this binding alters the structure and mobility of AF2, although there is no direct interaction between AF2 and TBP [56]. Since AF1 interact with both enhancer associating coactivators and promoter binding TBP, it is tempting to speculate that AF1 facilitate enhancer-promoter looping, in which enhancer is brought in contact with its target promoter through coactivators and general transcription factors [57]. This is also supported by the evidence that TATA-binding protein (TBP) binding to glucocorticoid receptor AF1 induces AF1 folding and facilitates its interaction with SRC-1 [20], which in turn may bring the promoter and enhancer together.

In summary, AF1 is important for PR to regulate cell growth adhesion and apoptosis in response to progestin. AF1 is involved in progestin regulation of two-thirds of PR target genes including genes involved in growth regulation, hypoxia, and TNFA Signalling via NFKB. We propose that AF1 monomethylation modulates PR-chromatin binding kinetics through modulating interaction with coregulators and general transcription factors. However, ligand-independent activity of AF1 operates on distinct mechanisms from that of ligand-dependent activities based on genome-wide gene expression analysis. Further experiments are required to demonstrate that AF1-FFF mutant indeed alters the kinetics of PR-chromatin interaction and enhancer-promoter looping. Importantly, as AF1 is required for the biphasic effect of progestin on cell proliferation in breast cancer cells, it may be targeted with small molecules to achieve desired therapeutic outcomes.

## Method and materials

### Cell culture

MCF-7 cell line was sourced from the American Type Culture Collection (ATCC). Cells were cultured in Dulbecco’s Modified Eagle’s Medium (DMEM) with phenol red (08488-55, Nacalai Tesque) supplemented with 7.5% fetal calf serum (FCS) and 2mM L-glutamine (Hyclone Logan, UT). Cultures were maintained at 37□ in a humidified incubator with 5% CO_2_ in air.

### Gene cloning

Site-directed mutagenesis was employed to generate PR AF1 K464F_K481F_R492F and K464Q_K481Q_R492Q mutants as previously reported [22]. PRB cDNA was cloned in pLVX-Puro vector using XhoI and Xbal restriction enzyme sites. Plasmid vectors carrying PRB and mutant cDNAs were designated as PRB, PRB-FFF and PRB-QQQ, respectively. An empty vector (EV) served as control. All constructs were verified by DNA sequencing.

### Generation of MCF-7 stable cell lines overexpressing PRB and PR AF1 mutants

MCF-7 cells were seeded at 1×10^5^ cells per well in a 6-well plate. 24 hours later, cells were transduced with lentiviruses carrying PRB or AF1 mutant cDNA at Multiplicity of Infection (MOI) of 2 in the presence of polybrene. The cells were then centrifuged for 90 minutes at 30°C and 1000xg with maximum acceleration and minimum deceleration without braking. Cells were then incubated for 48 hours before transferred to a 100-mm culture dish and cultured in selection medium containing 2.5ug/mL puromycin. A second selection was performed five days later after allowing the cells to recover for one week. The resulting stable MCF-7 cell lines expressing PRB, PRB-FFF and PRB-QQQ and empty vector EV were subsequently maintained in antibiotic free DMEM, and all experiments were conducted within 10 passages. Stable PR overexpression in mutant cell lines was routinely verified via western blotting and immunofluorescence.

### Hormone treatment

Cells were cultured for 48 hours in phenol-red-free DMEM supplemented with 5% dextran-charcoal-treated FCS (DCC-FCS) and 2mM L-glutamine prior to hormone treatment. Hormone treatment includes promegestone (R5020), estrogen (17-beta estradiol) or combination of both at designated concentrations in 0.01% ethanol in accordance with experimental design. Control groups received 0.01% ethanol alone.

### Cell Cycle analysis

Cells after 16 hours and 48 hours 10nM R5020 treatment were detached by 0.05% trypsin and stained with propidium iodide (PI) in Vindelov’s cocktail [10 mM Tris-HCl (pH 8), 10 mM NaCl, 50 mg PI/l, 10 mg/l RNaseA, and 0.1% NP40]. Staining was performed in dark for 30 minutes at 4°C. Samples were then analysed using BD LSRFortessa™ X-20 flow cytometer (BD Biosciences, Franklin Lakes, NJ, USA) with excitation at 488 nm. Cell cycle distribution was determined using Flowjo software.

### Cell imaging and fluorescent microscopy

Cells were seeded at a density of 1 × 10^5^ cells per 35-mm dish, each with a glass coverslip in phenol-free DMEM supplemented with 5% DCC-FCS and 2 mM L-glutamine. After 48 hours, cells were treated with either 0.01% ETOH (vehicle control) or 10 nM R5020. Cell images were captured using Olympus phase contrast microscope (XL7) after 24-, 48-, 72- and 96-hours post-treatment.

For PR immunostaining, transduced cells grown on glass coverslip were fixed with 3.7% formaldehyde and permeabilized with 0.2% Triton X-100, each for 10 minutes. Cells were then blocked for 1 hour in blocking agent (2% FBS in PBS), followed by incubation with anti-PR antibody H190 (Santa Cruz Biotechnology Inc., Dallas, TX, USA, LOT number sc-7208) in 1:200 dilution at 37 °C for 2 hours. After washing, cells were incubated with Alexa Fluor™ 488-conjugated Goat anti-Rabbit IgG with (Invitrogen, Carlsbad, California, USA, Cat # A-11034) for 1 hour. Coverslips were then mounted using Duolink^®^ mounting medium with DAPI (Cat # -82040-0005) for immunofluorescence imaging.

### Protein Lysate Collection and Western Blotting Analysis

Cells were lysed using cold lysis buffer containing 50 mM HEPES-KOH (pH 7.5), 100 mM NaF, 150mM NaCl and 1% Triton X-100. Lysates were collected from the supernatant after centrifugation at 12,000xg for 12 minutes. Proteins in total cell lysates were quantitated and resolved by SDS-PAGE electrophoresis and transferred onto a nitrocellulose or PVDF membrane. Membranes were blocked with 5% skimmed milk or 2.5% BSA in Tris buffered saline with Tween 20 (TBST), followed by overnight incubation with primary antibodies.

The primary antibodies used in the experiments are: H190 Total PR (Santa Cruz Biotechnology Inc., Dallas, TX, USA, LOT number sc-7208), ERα (Santa Cruz Biotechnology Inc., Dallas, TX, USA, LOT number sc-8002), BCL-2 (Santa Cruz Biotechnology Inc., Dallas, TX, USA, LOT number sc-7382), BNIP3 (Cell Signaling Technology, Danvers, MA, USA, #44060), BNIP3L/NIX (Cell Signaling Technology, Danvers, MA, USA, #12396), HTRA2/OMI (Cell Signaling Technology, Danvers, MA, USA, #9745), Endonuclease G (Cell Signaling Technology, Danvers, MA, USA, #4969) and GAPDH (Santa Cruz Biotechnology Inc., Dallas, TX, USA, LOT number sc-47724). Primary antibodies were used at dilution of 1:1000, except for GAPDH, which was diluted at 1:10000.

Horseradish peroxidase (HRP)-conjugated secondary antibodies, anti-mouse (Cell Signaling Technology, Danvers, MA, USA, #7076) and anti-rabbit (Cell Signaling Technology, Danvers, MA, USA, #7074) were used based on the host species of primary antibodies at dilution of 1:1000. Proteins detection was performed using Immobilon Western Chemiluminescent HRP substrate (Merck Millipore, Billerica, MA, USA) and ChemiDoc MP Imaging System (Bio-Rad Laboratories, Inc).

### RNA Extraction

Following hormone treatment, cells cultured in 60-mm dishes were lysed in cold TRIzol reagent, and total RNA was isolated according to the manufacturer’s protocol (Life Technologies, MD, USA). RNA was precipitated using isopropanol and pelleted at 12000xg, 4°C for 15 minutes. The pelleted RNA was washed twice with 75% ethanol prepared in DEPC-treated water, briefly air-dried and resuspended in DEPC-treated water.

Total RNA concentration and purity was accessed using a Nanodrop 2000 spectrophotometer (Thermo Fisher Scientific, USA). RNA integrity was further evaluated through agarose gel electrophoresis to check for RNA degradation.

### RNA-Seq analysis

Total RNA was treated with DNase I using the DNA-free™ DNA Removal Kit (Invitrogen, Carlsbad, CA, USA) to eliminate residual genomic DNA. DNA free RNA samples were submitted to Genome Institute of Singapore (Agency for Science, Technology and Research) for library preparation and paired-end sequencing using Illumina HiSeq4000 platform.

Sequencing read quality was assessed using *FastQC*. Transcription quantification was performed using the STAR aligners against the human reference genome GRCh38 (hg38) obtained from Ensembl. Transcript-level counts were summarized to gene-level counts using R package *tximport* [58]. Differential gene expression analysis was performed using *DESeq2* [59]. Genes with an adjusted p-value less than 0.05 (padj<0.05) were considered as differential expressed genes (DEGs). DESeq2-normalized counts data from were used for subsequent downstream analysis. Heatmaps of differentially expressed genes were generated using the online platform SRplot.

### Gene set enrichment analysis (GSEA)

GSEA was performed using the software downloaded from the official website https://www.gsea-msigdb.org/gsea/index.jsp. Analyses were conducted using default parameters. Enriched gene set with a false discovery rate (FDR) of q < 0.25 were considered as statistically significant. The Hallmark gene sets from the Molecular Signature Database (MSigDB) were used, which represents the curated gene sets characterizing well-defined biological processes [60, 61].

### ChIP-seq and ChIP-qPCR

Following treatment with vehicle or 10nM R5020 for 1 hour, cells were crosslinked with formaldehyde at a final concentration of 1% for 10 minutes at room temperature. The cross-linking was quenched by the addition of glycine to a final concentration of 0.125 M. Cells were then harvested, lysed, and subjected to sonication using Diagenode Bioruptor Pico at 5 cycles of 30s “ON” 30s “OFF”. After centrifugation to remove cell debris, 1% of lysate was reserved as input control.

Chromatin immunoprecipitation was performed using anti-PR antibody H-190 (Santa Cruz Biotechnology Inc., Dallas, TX, USA, LOT number sc-7208). The lysate was incubated with 5 ug anti-PR antibody overnight at 4 °C. Protein-chromatin complex was captured using Pierce™ ChIP-grade Protein A/G Magnetic Beads (ThermoFisher Scientific, Cat. #26162). Chromatin was eluted, and crosslinks were reversed by treatment with RNase A and Proteinase K. Input samples and ChIP’ed DNA was purified using QIAquick PCR Purification Kit (Qiagen). DNA concentrations were quantified using Qubit Fluorometer (ThermoFisher Scientific). The samples were submitted to Genome Institute of Singapore (Agency for Science, Technology and Research) for library preparation and paired-end sequencing using Illumina HiSeq4000 platform.

ChIP Quantitative real-time PCR was carried out using SYBR Green master mix (KAPA) on an ABI Prism 7700 sequence detection system (Applied Biosystem) based on the manufacturer’s protocol. Real-time PCR for each targeted gene was performed in duplicates. The fold enrichment of target sequence in the immunoprecipitated (IP) fraction was calculated using the comparative Ct method, normalized to the input fraction and expressed relative to vehicle treated controls. Primer sequence listed as followed [21, 62].

FKBP5 FW: 5’-TAATAGAGGGGCGAGAAGGCAGA-3’

FKBP5 RV: 5’-GGTAAGTGGGTGTGCTCGCTCA-3’

CHR6P FW: 5’-TCAGGAACAGTACACGAACGA-3’

CHR6P RV: 5’-CTGGCTCATCTTTCAGCACA-3’

### ChIP-seq Analysis

Model-based analysis of ChIP-seq version 2 (MACS2) was used to perform peak calling for ChIP-seq data, following default parameter [63]. Integrative Genome Viewer (IGV) was used to create representative snapshot for the peaks called [64]]. Diffbind, a R Bioconductor package was used to identify the peaks with differential binding between cell lines and treatment [65].

Binding sites were extended by 50 bp on both 5’ and 3’ ends, and the corresponding FASTA sequence were fetched using the Galaxy platform [66]. The sequences were scanned for motif matches using MEME-ChIP suite[67]. Motif analysis was also performed with the Seqpos motif tool [68].

### Statistical analysis

All statistical analysis was performed using the Graphpad Prism 9 software. Data are expressed as mean ± SEM (standard error of the mean). The degree of statistical significance is indicated with asterisks (* p< 0.05, ** p < 0.01, *** p < 0.001, **** p < 0.001).

## Supporting information

Supplementary Doc 1

Supplementary Doc 2

Supplementary Doc 3

Supplementary Doc 4

Supplementary Doc 5

## References

[1] J.D. Graham, C.L. Clarke, Physiological Action of Progesterone in Target Tissues*, Endocr Rev, 18 (1997) 502–519.

[2] J.J. Kim, E. Chapman-Davis, Role of progesterone in endometrial cancer, Semin Reprod Med, 28 (2010) 81–90.

[3] P.A. Joshi, H.W. Jackson, A.G. Beristain, M.A. Di Grappa, P.A. Mote, C.L. Clarke, J. Stingl, P.D. Waterhouse, R. Khokha, Progesterone induces adult mammary stem cell expansion, Nature, 465 (2010) 803–807.

[4] R.D. Rajaram, D. Buric, M. Caikovski, A. Ayyanan, J. Rougemont, J. Shan, S.J. Vainio, O. Yalcin-Ozuysal, C. Brisken, Progesterone and Wnt4 control mammary stem cells via myoepithelial crosstalk, Embo J, 34 (2015) 641–652.

[5] J.E. Rossouw, G.L. Anderson, R.L. Prentice, A.Z. LaCroix, C. Kooperberg, M.L. Stefanick, R.D. Jackson, S.A. Beresford, B.V. Howard, K.C. Johnson, J.M. Kotchen, J. Ockene, Risks and benefits of estrogen plus progestin in healthy postmenopausal women: principal results From the Women’s Health Initiative randomized controlled trial, Jama, 288 (2002) 321–333.

[6] V. Beral, Breast cancer and hormone-replacement therapy in the Million Women Study, Lancet, 362 (2003) 419–427.

[7] A.Z. Bluming, H.N. Hodis, R.D. Langer, ’Tis but a scratch: a critical review of the Women’s Health Initiative evidence associating menopausal hormone therapy with the risk of breast cancer, Menopause, 30 (2023) 1241–1245.

[8] H.N. Hodis, P.M. Sarrel, Menopausal hormone therapy and breast cancer: what is the evidence from randomized trials?, Climacteric, 21 (2018) 521–528.

[9] G.L. Anderson, R.T. Chlebowski, J.E. Rossouw, R.J. Rodabough, A. McTiernan, K.L. Margolis, A. Aggerwal, J. David Curb, S.L. Hendrix, F. Allan Hubbell, J. Khandekar, D.S. Lane, N. Lasser, A.M. Lopez, J. Potter, C. Ritenbaugh, Prior hormone therapy and breast cancer risk in the Women’s Health Initiative randomized trial of estrogen plus progestin, Maturitas, 55 (2006) 103–115.

[10] J.E. Manson, R.T. Chlebowski, M.L. Stefanick, A.K. Aragaki, J.E. Rossouw, R.L. Prentice, G. Anderson, B.V. Howard, C.A. Thomson, A.Z. LaCroix, J. Wactawski-Wende, R.D. Jackson, M. Limacher, K.L. Margolis, S. Wassertheil-Smoller, S.A. Beresford, J.A. Cauley, C.B. Eaton, M. Gass, J. Hsia, K.C. Johnson, C. Kooperberg, L.H. Kuller, C.E. Lewis, S. Liu, L.W. Martin, J.K. Ockene, M.J. O’Sullivan, L.H. Powell, M.S. Simon, L. Van Horn, M.Z. Vitolins, R.B. Wallace, Menopausal hormone therapy and health outcomes during the intervention and extended poststopping phases of the Women’s Health Initiative randomized trials, Jama, 310 (2013) 1353–1368.

[11] S. Giulianelli, J.P. Vaque, R. Soldati, V. Wargon, S.I. Vanzulli, R. Martins, E. Zeitlin, A.A. Molinolo, L.A. Helguero, C.A. Lamb, J.S. Gutkind, C. Lanari, Estrogen receptor alpha mediates progestin-induced mammary tumor growth by interacting with progesterone receptors at the cyclin D1/MYC promoters, Cancer Res, 72 (2012) 2416–2427.

[12] H. Singhal, M.E. Greene, G. Tarulli, A.L. Zarnke, R.J. Bourgo, M. Laine, Y.-F. Chang, S. Ma, A.G. Dembo, G.V. Raj, T.E. Hickey, W.D. Tilley, G.L. Greene, Genomic agonism and phenotypic antagonism between estrogen and progesterone receptors in breast cancer, Science Advances, 2 (2016) e1501924.

[13] H. Singhal, M.E. Greene, A.L. Zarnke, M. Laine, R. Al Abosy, Y.F. Chang, A.G. Dembo, K. Schoenfelt, R. Vadhi, X. Qiu, P. Rao, B. Santhamma, H.B. Nair, K.J. Nickisch, H.W. Long, L. Becker, M. Brown, G.L. Greene, Progesterone receptor isoforms, agonists and antagonists differentially reprogram estrogen signaling, Oncotarget, 9 (2018) 4282–4300.

[14] H. Mohammed, I.A. Russell, R. Stark, O.M. Rueda, T.E. Hickey, G.A. Tarulli, A.A. Serandour, S.N. Birrell, A. Bruna, A. Saadi, S. Menon, J. Hadfield, M. Pugh, G.V. Raj, G.D. Brown, C. D’Santos, J.L.L. Robinson, G. Silva, R. Launchbury, C.M. Perou, J. Stingl, C. Caldas, W.D. Tilley, J.S. Carroll, Progesterone receptor modulates ERα action in breast cancer, Nature, 523 (2015) 313–317.

[15] K.K. Hill, S.C. Roemer, M.E.A. Churchill, D.P. Edwards, Structural and Functional Analysis of Domains of the Progesterone Receptor, Mol Cell Endocrinol, 348 (2012) 418–429.

[16] L. Tora, H. Gronemeyer, B. Turcotte, M.P. Gaub, P. Chambon, The N-Terminal Region of the Chicken Progesterone-Receptor Specifies Target Gene Activation, Nature, 333 (1988) 185–188.

[17] L. Tora, J. White, C. Brou, D. Tasset, N. Webster, E. Scheer, P. Chambon, The human estrogen receptor has two independent nonacidic transcriptional activation functions, Cell, 59 (1989) 477–487.

[18] H. Jafari, S. Hussain, M.J. Campbell, Nuclear Receptor Coregulators in Hormone-Dependent Cancers, Cancers (Basel), 14 (2022).

[19] D.M. Lonard, B.W. O’Malley, Nuclear receptor coregulators: modulators of pathology and therapeutic targets, Nat Rev Endocrinol, 8 (2012) 598–604.

[20] S.H. Khan, S. Awasthi, C. Guo, D. Goswami, J. Ling, P.R. Griffin, S.S. Simons, Jr., R. Kumar, Binding of the N-terminal region of coactivator TIF2 to the intrinsically disordered AF1 domain of the glucocorticoid receptor is accompanied by conformational reorganizations, J Biol Chem, 287 (2012) 44546–44560.

[21] H.H. Chung, S.K. Sze, A.S. Tay, V.C. Lin, Acetylation at lysine 183 of progesterone receptor by p300 accelerates DNA binding kinetics and transactivation of direct target genes, J Biol Chem, 289 (2014) 2180–2194.

[22] A.R.E. Woo, S.K. Sze, H.H. Chung, V.C. Lin, Delineation of critical amino acids in activation function 1 of progesterone receptor for recruitment of transcription coregulators, Biochim Biophys Acta Gene Regul Mech, 1862 (2019) 522–533.

[23] S.H. Lee, C.L. Lim, W. Shen, S.M.X. Tan, A.R.E. Woo, Y.H.Y. Yap, C.A.S. Sian, W.W.B. Goh, W.-P. Yu, L. Li, V.C.L. Lin, Activation function 1 of progesterone receptor is required for progesterone antagonism of oestrogen action in the uterus, BMC Biology, 20 (2022) 222.

[24] S.H. Lee, C.L. Lim, W. Shen, S.M.X. Tan, A.R.E. Woo, Y.H.Y. Yap, C.A.S. Sian, W.W.B. Goh, W.P. Yu, L. Li, V.C.L. Lin, Activation function 1 of progesterone receptor is required for progesterone antagonism of oestrogen action in the uterus, BMC Biol, 20 (2022) 222.

[25] R.J. Andersen, N.R. Mawji, J. Wang, G. Wang, S. Haile, J.K. Myung, K. Watt, T. Tam, Y.C. Yang, C.A. Banuelos, D.E. Williams, I.J. McEwan, Y. Wang, M.D. Sadar, Regression of castrate-recurrent prostate cancer by a small-molecule inhibitor of the amino-terminus domain of the androgen receptor, Cancer Cell, 17 (2010) 535–546.

[26] T.T. Tran, C.H. Song, K.J. Kim, K. Lee, A new compound targets the AF-1 of androgen receptor and decreases its activity and protein levels in prostate cancer cells, Am J Cancer Res, 10 (2020) 4607–4623.

[27] J. Zhu, X. Salvatella, P. Robustelli, Small molecules targeting the disordered transactivation domain of the androgen receptor induce the formation of collapsed helical states, Nat Commun, 13 (2022) 6390.

[28] S.D. Groshong, G.I. Owen, B. Grimison, I.E. Schauer, M.C. Todd, T.A. Langan, R.A. Sclafani, C.A. Lange, K.B. Horwitz, Biphasic regulation of breast cancer cell growth by progesterone: role of the cyclin-dependent kinase inhibitors, p21 and p27(Kip1), Mol Endocrinol, 11 (1997) 1593–1607.

[29] E.A. Musgrove, C.S.L. Lee, R.L. Sutherland, Progestins Both Stimulate and Inhibit Breast-Cancer Cell-Cycle Progression While Increasing Expression of Transforming Growth Factor-Alpha, Epidermal Growth-Factor Receptor, C-Fos, and C-Myc Genes, Mol Cell Biol, 11 (1991) 5032–5043.

[30] N. Bajalovic, Y.Z. Or, A.R.E. Woo, S.H. Lee, V.C.L. Lin, High Levels of Progesterone Receptor B in MCF-7 Cells Enable Radical Anti-Tumoral and Anti-Estrogenic Effect of Progestin, Biomedicines, 10 (2022).

[31] Y. Suzuki, Y. Imai, H. Nakayama, K. Takahashi, K. Takio, R. Takahashi, A serine protease, HtrA2, is released from the mitochondria and interacts with XIAP, inducing cell death, Mol Cell, 8 (2001) 613–621.

[32] L.Y. Li, X. Luo, X. Wang, Endonuclease G is an apoptotic DNase when released from mitochondria, Nature, 412 (2001) 95–99.

[33] E.J. Faivre, C.A. and Lange, Progesterone Receptors Upregulate Wnt-1 To Induce Epidermal Growth Factor Receptor Transactivation and c-Src-Dependent Sustained Activation of Erk1/2 Mitogen-Activated Protein Kinase in Breast Cancer Cells, Mol Cell Biol, 27 (2007) 466–480.

[34] C.R. Hagan, T.M. Regan, G.E. Dressing, C.A. Lange, ck2-dependent phosphorylation of progesterone receptors (PR) on Ser81 regulates PR-B isoform-specific target gene expression in breast cancer cells, Mol Cell Biol, 31 (2011) 2439–2452.

[35] D.B. Hardy, B.A. Janowski, C.-C. Chen, C.R. Mendelson, Progesterone Receptor Inhibits Aromatase and Inflammatory Response Pathways in Breast Cancer Cells via Ligand-Dependent and Ligand-Independent Mechanisms, Mol Endocrinol, 22 (2008) 1812–1824.

[36] L.Q.J. Lim, L. Adler, E. Hajaj, L.R. Soria, R.B. Perry, N. Darzi, R. Brody, N. Furth, M. Lichtenstein, E. Bab-Dinitz, Z. Porat, T. Melman, A. Brandis, S. Malitsky, M. Itkin, Y. Aylon, S. Ben-Dor, I. Orr, A. Pri-Or, R. Seger, Y. Shaul, E. Ruppin, M. Oren, M. Perez, J. Meier, N. Brunetti-Pierri, E. Shema, I. Ulitsky, A. Erez, ASS1 metabolically contributes to the nuclear and cytosolic p53-mediated DNA damage response, Nat Metab, 6 (2024) 1294–1309.

[37] W. Luo, Z. Zou, Y. Nie, J. Luo, Z. Ming, X. Hu, T. Luo, M. Ouyang, M. Liu, H. Tang, Y. Xie, K. Peng, L. Chen, J. Zhou, Z. Luo, ASS1 inhibits triple-negative breast cancer by regulating PHGDH stability and de novo serine synthesis, Cell Death Dis, 15 (2024) 319.

[38] Y. Wang, K. Li, W. Zhao, Z. Liu, J. Liu, A. Shi, T. Chen, W. Mu, Y. Xu, C. Pan, Z. Zhang, Aldehyde dehydrogenase 3B2 promotes the proliferation and invasion of cholangiocarcinoma by increasing Integrin Beta 1 expression, Cell Death Dis, 12 (2021) 1158.

[39] B. Jaeger, J.C. Schupp, L. Plappert, O. Terwolbeck, N. Artysh, G. Kayser, P. Engelhard, T.S. Adams, R. Zweigerdt, H. Kempf, S. Lienenklaus, W. Garrels, I. Nazarenko, D. Jonigk, M. Wygrecka, D. Klatt, A. Schambach, N. Kaminski, A. Prasse, Airway basal cells show a dedifferentiated KRT17(high)Phenotype and promote fibrosis in idiopathic pulmonary fibrosis, Nat Commun, 13 (2022) 5637.

[40] P. Yin, D. Roqueiro, L. Huang, J.K. Owen, A. Xie, A. Navarro, D. Monsivais, J.S.C. V, J.J. Kim, Y. Dai, S.E. Bulun, Genome-Wide Progesterone Receptor Binding: Cell Type-Specific and Shared Mechanisms in T47D Breast Cancer Cells and Primary Leiomyoma Cells, PLoS One, 7 (2012) e29021.

[41] H. Singhal, M.E. Greene, G. Tarulli, A.L. Zarnke, R.J. Bourgo, M. Laine, Y.F. Chang, S. Ma, A.G. Dembo, G.V. Raj, T.E. Hickey, W.D. Tilley, G.L. Greene, Genomic agonism and phenotypic antagonism between estrogen and progesterone receptors in breast cancer, Sci Adv, 2 (2016) e1501924.

[42] J.C. Leo, V.C. Lin, The activities of progesterone receptor isoform A and B are differentially modulated by their ligands in a gene-selective manner, Int J Cancer, 122 (2008) 230–243.

[43] M. Singh, V.R. Krishnamoorthy, S. Kim, S. Khurana, H.M. LaPorte, Brain-derived neuerotrophic factor and related mechanisms that mediate and influence progesterone-induced neuroprotection, Front Endocrinol (Lausanne), 15 (2024) 1286066.

[44] G.M. Bernardo, K.L. Lozada, J.D. Miedler, G. Harburg, S.C. Hewitt, J.D. Mosley, A.K. Godwin, K.S. Korach, J.E. Visvader, K.H. Kaestner, F.W. Abdul-Karim, M.M. Montano, R.A. Keri, FOXA1 is an essential determinant of ER alpha expression and mammary ductal morphogenesis, Development, 137 (2010) 2045–2054.

[45] C.L. Clarke, J.D. Graham, Non-overlapping progesterone receptor cistromes contribute to cell-specific transcriptional outcomes, PLoS One, 7 (2012) e35859.

[46] N.L. Weigel, Y. Zhang, Ligand-independent activation of steroid hormone receptors, J Mol Med, 76 (1998) 469–479.

[47] R. White, M. Sjoberg, E. Kalkhoven, M.G. Parker, Ligand-independent activation of the oestrogen receptor by mutation of a conserved tyrosine, Embo J, 16 (1997) 1427–1435.

[48] Y. Sun, H.H. Chung, A.R. Woo, V.C. Lin, Protein arginine methyltransferase 6 enhances ligand-dependent and -independent activity of estrogen receptor alpha via distinct mechanisms, Biochim Biophys Acta, 1843 (2014) 2067–2078.

[49] G.L. Hager, J.G. McNally, T. Misteli, Transcription dynamics, Mol Cell, 35 (2009) 741–753.

[50] W.N. Jefferson, T. Wang, E. Padilla-Banks, C.J. Williams, Unexpected nuclear hormone receptor and chromatin dynamics regulate estrous cycle dependent gene expression, Nucleic Acids Res, 52 (2024) 10897–10917.

[51] C. Carlberg, S. Seuter, Dynamics of nuclear receptor target gene regulation, Chromosoma, 119 (2010) 479–484.

[52] R.I. Morimoto, Dynamic remodeling of transcription complexes by molecular chaperones, Cell, 110 (2002) 281–284.

[53] Z. Liu, Y. Chen, Q. Xia, M. Liu, H. Xu, Y. Chi, Y. Deng, D. Xing, Linking genome structures to functions by simultaneous single-cell Hi-C and RNA-seq, Science, 380 (2023) 1070–1076.

[54] H.H. Chung, S.K. Sze, A.R. Woo, Y. Sun, K.H. Sim, X.M. Dong, V.C. Lin, Lysine methylation of progesterone receptor at activation function 1 regulates both ligand-independent activity and ligand sensitivity of the receptor, J Biol Chem, 289 (2014) 5704–5722.

[55] X. Yu, P. Yi, A.K. Panigrahi, L.E.V. Lumahan, J.P. Lydon, D.M. Lonard, S.J. Lutdke, Z. Wang, B.W. O’Malley, Spatial definition of the human progesterone receptor-B transcriptional complex, iScience, 25 (2022) 105321.

[56] D. Goswami, C. Callaway, B.D. Pascal, R. Kumar, D.P. Edwards, P.R. Griffin, Influence of domain interactions on conformational mobility of the progesterone receptor detected by hydrogen/deuterium exchange mass spectrometry, Structure, 22 (2014) 961–973.

[57] M.J. Friedman, T. Wagner, H. Lee, M.G. Rosenfeld, S. Oh, Enhancer-promoter specificity in gene transcription: molecular mechanisms and disease associations, Exp Mol Med, 56 (2024) 772–787.

[58] C. Soneson, M.I. Love, M.D. Robinson, Differential analyses for RNA-seq: transcript-level estimates improve gene-level inferences, F1000Res, 4 (2015) 1521.

[59] M.I. Love, W. Huber, S. Anders, Moderated estimation of fold change and dispersion for RNA-seq data with DESeq2, Genome Biol, 15 (2014) 550.

[60] V.K. Mootha, C.M. Lindgren, K.F. Eriksson, A. Subramanian, S. Sihag, J. Lehar, P. Puigserver, E. Carlsson, M. Ridderstrale, E. Laurila, N. Houstis, M.J. Daly, N. Patterson, J.P. Mesirov, T.R. Golub, P. Tamayo, B. Spiegelman, E.S. Lander, J.N. Hirschhorn, D. Altshuler, L.C. Groop, PGC-1alpha-responsive genes involved in oxidative phosphorylation are coordinately downregulated in human diabetes, Nat Genet, 34 (2003) 267–273.

[61] A. Subramanian, P. Tamayo, V.K. Mootha, S. Mukherjee, B.L. Ebert, M.A. Gillette, A. Paulovich, S.L. Pomeroy, T.R. Golub, E.S. Lander, J.P. Mesirov, Gene set enrichment analysis: a knowledge-based approach for interpreting genome-wide expression profiles, Proc Natl Acad Sci U S A, 102 (2005) 15545–15550.

[62] R. Zaurin, R. Ferrari, A.S. Nacht, J. Carbonell, F. Le Dily, J. Font-Mateu, L.I. de Llobet Cucalon, E. Vidal, A. Lioutas, M. Beato, G.P. Vicent, A set of accessible enhancers enables the initial response of breast cancer cells to physiological progestin concentrations, Nucleic Acids Res, 49 (2021) 12716–12731.

[63] Y. Zhang, T. Liu, C.A. Meyer, J. Eeckhoute, D.S. Johnson, B.E. Bernstein, C. Nusbaum, R.M. Myers, M. Brown, W. Li, X.S. Liu, Model-based analysis of ChIP-Seq (MACS), Genome Biol, 9 (2008) R137.

[64] J.T. Robinson, H. Thorvaldsdottir, W. Winckler, M. Guttman, E.S. Lander, G. Getz, J.P. Mesirov, Integrative genomics viewer, Nat Biotechnol, 29 (2011) 24–26.

[65] C. Boeckel, X. Pastor, M. Heinig, T. Walzthoeni, Differential Analysis of Protein-DNA Binding Using ChIP-Seq Data, Methods Mol Biol, 2846 (2024) 63–89.

[66] C. Galaxy, The Galaxy platform for accessible, reproducible and collaborative biomedical analyses: 2022 update, Nucleic Acids Res, 50 (2022) W345–W351.

[67] T.L. Bailey, J. Johnson, C.E. Grant, W.S. Noble, The MEME Suite, Nucleic Acids Res, 43 (2015) W39–49.

[68] T. Liu, J.A. Ortiz, L. Taing, C.A. Meyer, B. Lee, Y. Zhang, H. Shin, S.S. Wong, J. Ma, Y. Lei, U.J. Pape, M. Poidinger, Y. Chen, K. Yeung, M. Brown, Y. Turpaz, X.S. Liu, Cistrome: an integrative platform for transcriptional regulation studies, Genome Biol, 12 (2011) R83.

