## Supplementary Figures for "Methylation mimic mutations of progesterone receptor AF1 impair gene-specific regulation through stabilized chromatin interactions"

### Supplementary Fig. 1

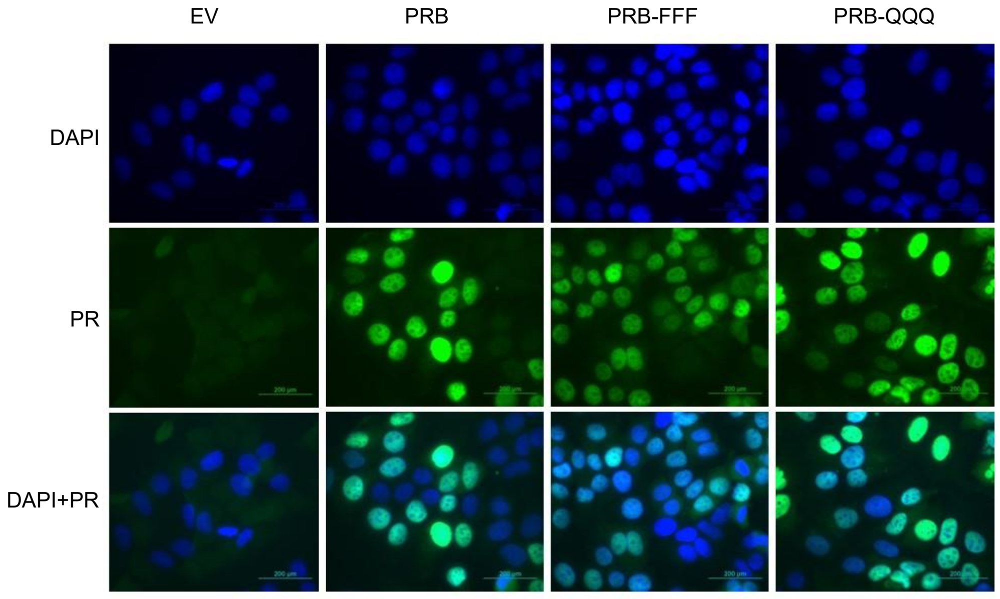


**Supplementary Fig. 1 PR AF1 mutants exhibit normal nuclear localization.** Immunofluorescence analysis verifying PR overexpression in transduced MCF-7 stable cell line. PR (green) shows nuclear localization. Nuclei are counterstained with DAPI (blue); merged images show DAPI+PR.

### Supplementary Fig. 2

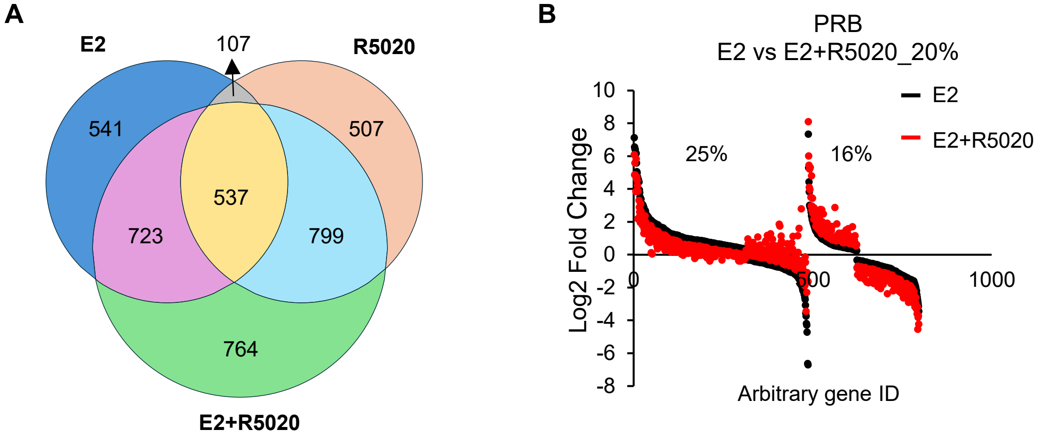


**Supplementary Fig. 2. AF1 modulated ligand independent activity of PR. (A)** Venn diagram indicates the overlap of genes significantly regulated by E2, R5020 and E2+R5020 in PRB cells (padj < 0.05). **(B)** 41% genes in PRB cells show more than 20% difference in regulation when treated with E2+R5020 compared to E2 alone. 25% of these shows less regulation by PRB.

### Supplementary Fig. 3

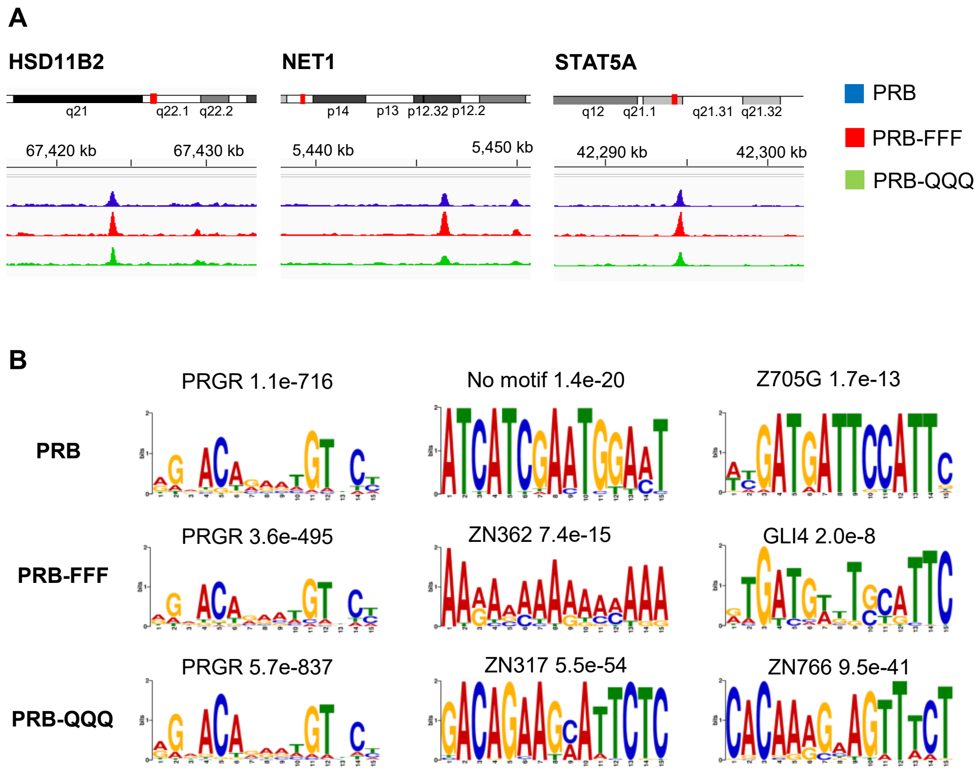


**Supplementary Fig. 3 Genome wide distribution of PR binding in PRB and AF1 mutants. (A)** Genome browser tracks showing ChIP-seq enrichment at representative PR target genes *Hsd11b2*, *Net1* and *Stat5a*. **(B)** The top 3 most significant binding motifs identified using MEME software are shown for PRB, PRB-FFF and PRB-QQQ cells. Enriched sequence with no match to a known transcription factor binding motif is indicated as “No motif”.
